# Environmental DNA (eDNA) metabarcoding differentiates between micro-habitats within the rocky intertidal

**DOI:** 10.1101/2023.08.03.551543

**Authors:** Meghan M. Shea, Alexandria B. Boehm

## Abstract

While the utility of environmental DNA (eDNA) metabarcoding surveys for biodiversity monitoring continues to be demonstrated, the spatial and temporal variability of eDNA, and thus the limits of the differentiability of an eDNA signal, remains under-characterized. In this study, we collected eDNA samples from distinct micro-habitats (∼40 m apart) in a rocky intertidal ecosystem over their exposure period in a tidal cycle. During this period, the micro-habitats transitioned from being interconnected, to physically isolated, to interconnected again. Using a well-established eukaryotic (cytochrome oxidase subunit I) metabarcoding assay, we detected 415 species across 28 phyla. Across a variety of univariate and multivariate analyses, using just taxonomically-assigned data as well as all detected amplicon sequence variants (ASVs), we identified unique eDNA signals from the different micro-habitats sampled. This differentiability paralleled ecological expectations and increased as the sites became more physically disconnected. Our results demonstrate that eDNA biomonitoring can differentiate micro-habitats in the rocky intertidal only 40 m apart, that these differences are ecologically meaningful, and that physical connectivity informs the degree of differentiation possible. These findings showcase the potential power of eDNA biomonitoring to increase the spatial and temporal resolution of marine biodiversity data, aiding research, conservation, and management efforts.

## 1. Introduction

Human impacts on marine ecosystems are substantial, yet heterogeneous—making it challenging to study human-induced changes at the scales necessary to address them (Halpern et al., 2019). In particular, coastal ecosystems experience both high cumulative impact as well as the fastest pace of increase in that impact across marine environments (Halpern et al., 2019). Yet biomonitoring of marine habitats is often limited. Many typical approaches, such as baited remote underwater video, transects, or diver tows, require particular skills and taxonomic expertise, suitable field conditions, and significant resources (Stat et al., 2017). To address these biomonitoring gaps, many have turned to environmental DNA (eDNA): genetic material found in environmental samples such as soil, sediment, water, and air (Taberlet et al., 2018). Intracellular and extracellular DNA—mainly originating from urine and fecal matter, shed epithelial cells, and tissue from decomposing organisms (Harrison et al., 2019)—extracted from these environmental matrices can be amplified and compared to known reference sequences to uncover what organisms might have contributed the collected DNA (Taberlet et al., 2018). Biomonitoring using eDNA can help address challenges with conventional biodiversity monitoring approaches, potentially lowering costs (Evans et al., 2017; Hering et al., 2018), enabling less invasive sampling (Beja-Pereira et al., 2009), and reducing the need for in-depth taxonomic knowledge during field sampling (Hering et al., 2018). Additionally, eDNA surveys have been validated as comparable, or complementary, to species identification using conventional morpho-taxonomic approaches (Fediajevaite et al., 2021; Keck et al., 2022; McElroy et al., 2020). For these reasons, eDNA biomonitoring has been used to answer a wide variety of research questions in marine environments (e.g. Berry et al., 2019; Gold, Sprague, et al., 2021; Stat et al., 2017; West et al., 2020), and has been increasingly positioned as a helpful tool for management (Gold, Sprague, et al., 2021; Gold et al., 2022; Kelly et al., 2014).

However a core difference between eDNA biomonitoring and conventional biodiversity monitoring approaches can complicate increased reliance on eDNA tools: the presence of DNA at a particular time and location does not necessarily correlate with a corresponding organism occurrence at the same spatial and temporal scale (Harrison et al., 2019). The concentration of eDNA in the environment is a function of the sources of that eDNA, the fate processes (e.g. decay) that remove it from the water column, and the transport processes that move it and dilute it (Harrison et al., 2019). These three processes are influenced by a range of biotic (e.g. organism life history characteristics, microbial communities) and abiotic (e.g. temperature, currents, sunlight) factors acting together in complex and environmentally-specific ways (Harrison et al., 2019). While there have been many efforts to study processes that modulate eDNA concentrations and presence in the environment (see Harrison et al., 2019), it is more challenging to then determine, given the sources, fate, and transport of eDNA in a particular environment, what scales of spatial and temporal differences are possible to detect.

In marine environments, several studies have sought to characterize the temporal and spatial variability of eDNA. While marine mesocosm experiments have indicated that fish eDNA can persist on the order of days (Minamoto et al., 2017; Sassoubre et al., 2016; Thomsen et al., 2012), field experiments demonstrate much shorter eDNA persistence; studies have shown that foreign eDNA introduced in marine environments was only detectable on the order of hours after being released (Ely et al., 2021; Murakami et al., 2019). Field studies have documented that eDNA can reflect changes in multivariate community structure over seasonal (Jia et al., 2020; Sevellec et al., 2020; Djurhuus et al., 2020) to sub-daily (Suter et al., 2020) time scales. Additionally, field studies have shown that eDNA can be spatially localized, with introduced foreign eDNA only detectable 10s of meters from its source (Ely et al., 2021; Murakami et al., 2019), and significant differences in species composition reflected in eDNA sampled from marine environments on the scale of meters vertically (Jeunen et al., 2020) and 10s of meters horizontally (Gold, Sprague, et al., 2021; O’Donnell et al., 2017; Port et al., 2016).

These experimental and descriptive studies of fate and transport of eDNA in marine environments highlight several key themes. For one, under certain conditions, eDNA can be considered quite spatially and temporally localized in marine environments—that is, an eDNA signal could be interpreted as representing primarily organisms present on the order of hours and 10s of meters from the sampling event. At the same time, these studies reinforce that the processes that influence eDNA concentrations interact in variable, site-specific ways that are hard to predict mechanistically—motivating a need to study the spatiotemporal dynamics of eDNA in marine environments that may differ substantially from those previously studied, such as rocky intertidal ecosystems.

Rocky intertidal ecosystems, some of the most physically variable and biodiverse marine environments on the planet (Helmuth et al., 2006), provide an important extension to previous work on the spatiotemporal variability of eDNA. For one, they are environments where ecological changes occur over particularly small spatial and temporal scales—across gradients of thermal and desiccation stresses, and across the tidal cycle—testing the limits of the differentiability of eDNA signals (Helmuth et al., 2006). Additionally, unlike previous experimental (Ely et al., 2021; Murakami et al., 2019) and modelling (Andruszkiewicz et al., 2019) studies which showed significant transport of eDNA in well-connected marine environments, rocky intertidal zones can experience large changes in physical connectivity; within a span of several hours, a site can transition from a well-mixed surf zone to a quiescent collection of isolated tide pools. Several studies have demonstrated that eDNA sampling in nearshore environments yields similar community composition across tides, suggesting that there is not a significant tidally-driven influx of offshore eDNA (Kelly et al., 2018; Lafferty et al., 2021). However, at smaller spatial and temporal scales, it is not clear how rapid changes in physical connectivity influence of transport of eDNA within the intertidal (Figure 1). For example, at high tide (Figure 1A), shedding, decay, and transport may all influence the eDNA signature from a particular micro-habitat. However, at low tide (Figure 1B), transport could be significantly diminished, potentially making it possible to detect a unique eDNA signature from that micro-habitat.

**Figure 1.**
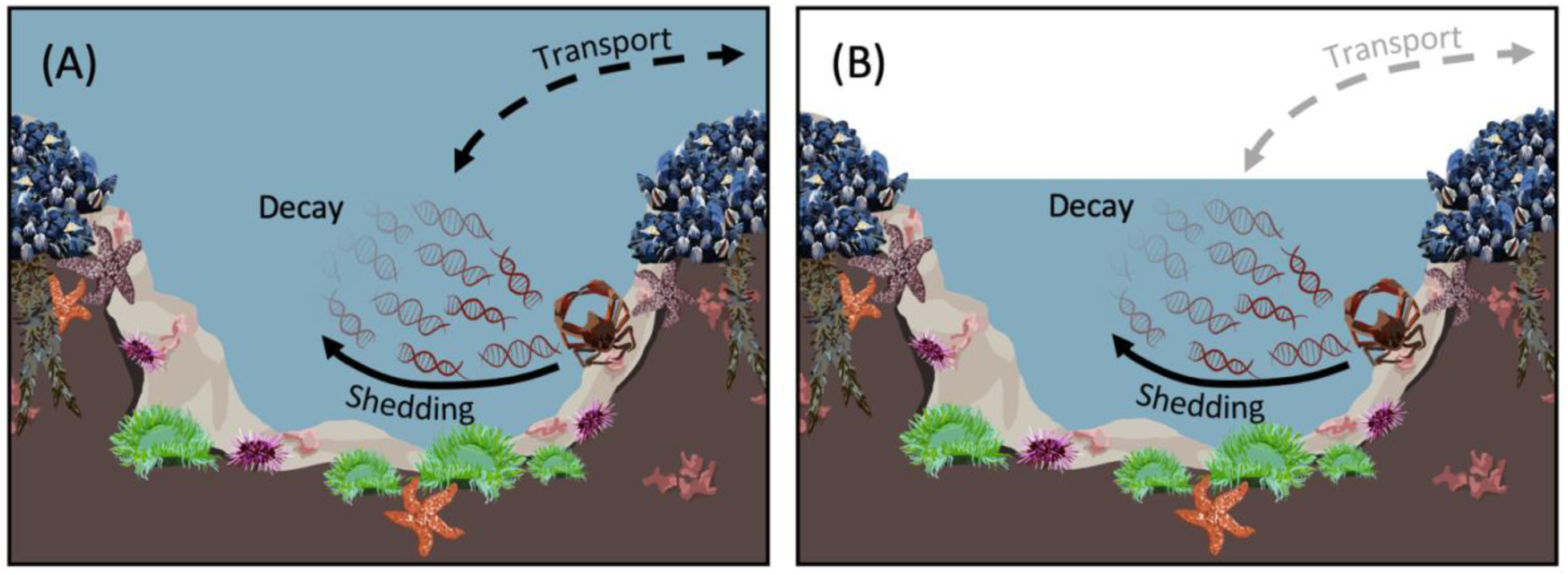
Conceptual model detailing the change in influences on the detectability of eDNA as physical connectivity in the intertidal changes between high tide (A) and low tide (B).

In this study, we use a rocky intertidal ecosystem to further characterize the spatiotemporal variability of eDNA in nearshore marine environments. Specifically, we sought to answer the following question: Does biodiversity detected using an eDNA metabarcoding survey differ between distinct micro-habitats in the rocky intertidal that vary in connectedness during the tidal cycle? We use the term biodiversity broadly to encompass multiple intersecting types and scales of analyses. We are interested in both unique eDNA sequences detected (amplicon sequence variants, or ASVs) independent of taxonomic information, as well as taxa detected, meaning the agglomeration of ASVs for which there is a species-level taxonomic assignment. Additionally, we are interested in both univariate changes in detection of individual ASVs and taxa, as well as multivariate changes in community composition. By distinct micro-habitats, we mean locations within 10s of meters of each other that are situated in different zones of the intertidal; substantial prior research using conventional ecological approaches has characterized that vertical and horizontal zonation in rocky intertidal shores informs community composition, so these micro-habitats would be expected to have different ecological communities (Connell, 1972). Our null hypothesis is that despite assumed underlying ecological differences, there are no changes in biodiversity detected via eDNA between micro-habitats given their proximity. However, we expect that eDNA metabarcoding will reflect both univariate and multivariate differences in biodiversity between micro-habitats, and that these differences will increase as the sites become physically isolated.

## 2. Materials and Methods

### 2.1. Reproducibility

To improve reproducibility (e.g. Dickie et al., 2018; Shea et al., 2023), enable open data science (e.g. Fredston & Lowndes, 2024), and aid in the initiation of new eDNA biomonitoring projects, we have published detailed, step-by-step protocols for many of the methods described, including specific materials used, photographs, and additional methodological notes not possible to include here. See Shea and Boehm for sample collection and filtering (2023b), for DNA extractions (2023c), for PCR amplification (2023d), and for shipping samples (2023e). Additionally, we have published all data and code for replicating our analyses via Dryad (Private Reviewer Link: https://datadryad.org/stash/share/Kxkudmvtnl8nEBBi3NBo2VFxFeluDLh1WlhmgR66lR0), including FASTQ files and eDNA datasets (pre-processing & post-processing). Through Dryad and Zenodo, our modified Anacapa Container and scripts for bioinformatics (Shea & Boehm, 2023a), as well as a GitHub repository including an R Markdown file that reproduces all methods & results detailed here (Shea & Boehm, 2023f) can also be accessed.

### 2.2. Sampling Site

To better understand spatial and temporal differences in eDNA signals in a complex coastal environment, we sought a rocky intertidal field location that had consistent, large, accessible tide pools that were fully isolated from one another at some low tides but interconnected during other parts of their exposure period. We selected the intertidal at Pillar Point, a headlands promontory to the west of Pillar Point Harbor in San Mateo County, California, USA. Pillar Point is a popular recreational intertidal site that is directly adjacent to the Pillar Point State Marine Conservation Area (no specific use scientific collection permit required).

Within Pillar Point, we sampled at three discrete locations: two individual tide pools with a range of physical connectivity (Tide Pool 1, S1: 37.495306°, -122.498744°; Tide Pool 2, S2: 37.494992°, -122.498955°) and an equidistant location (Nearshore, N: 37.495288°, -122.499198°) where there was well-mixed offshore water for the duration of the tidal cycle (Figure 2). Each site was approximately 40 meters from all other sites. Tide Pool 1 and Tide Pool 2 are fully isolated at tidal heights of around 0 m (mean low low water, MLLW) or lower, and substantially connected at around 0.25 m or higher. On the day we sampled, this meant water actively flowed between the locations at the start (11:30 PST) and end (17:00 PST) of the sampling period, but that the sites were disconnected at low tide in the middle of the sampling period.

**Figure 2.**
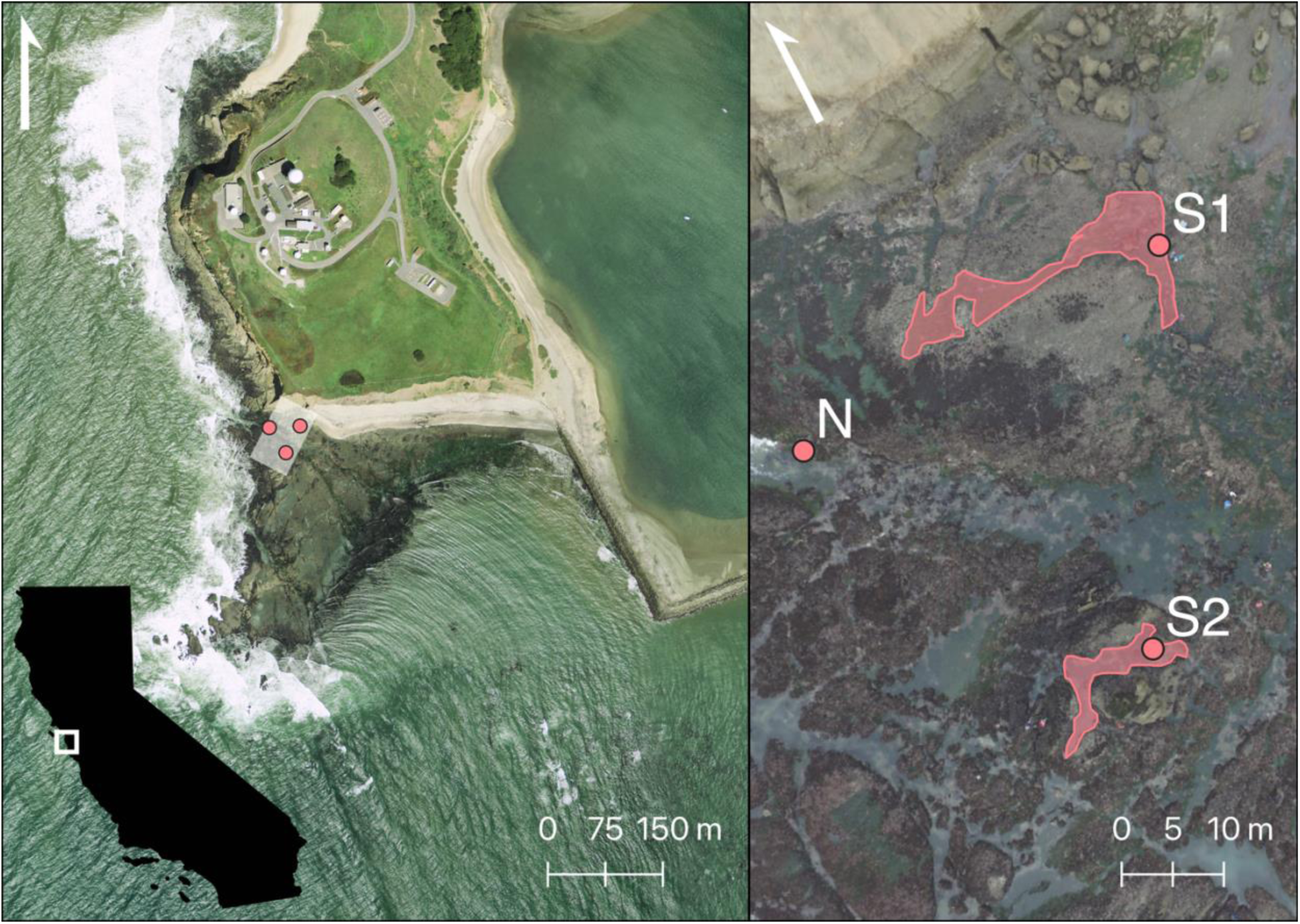
Depiction of sampling location using USGS high resolution orthoimagery (9 April 2011; left) and drone imagery (28 January 2022; right). The three individual sampling sites are marked and labeled, with polygons showing the approximate extant of tide pool locations S1 and S2 at low tide.

### 2.3. Sample Collection & Filtration

We collected 1 L surface samples from each location every 30 minutes for the duration of time the rocky intertidal was exposed on 28 January 2022, using single-use enteral feeding pouches (Covidien, Dublin, Ireland). Sampling commenced at 11:30 PST; the exact times of sample collection, in relation to the tide, are shown in Figure 3. Following the approach used by Gold et al. (2021), we attached a sterile 0.22 μm pore size Sterivex cartridge (MilliporeSigma, Burlington, MA, USA) to the tubing of each feeding pouch, allowing samples to be immediately gravity filtered in the field. While gravity filtering (1-2 hours per sample), samples were shaded with an awning to prevent any degradation by sunlight (Andruszkiewicz et al., 2017).

**Figure 3.**
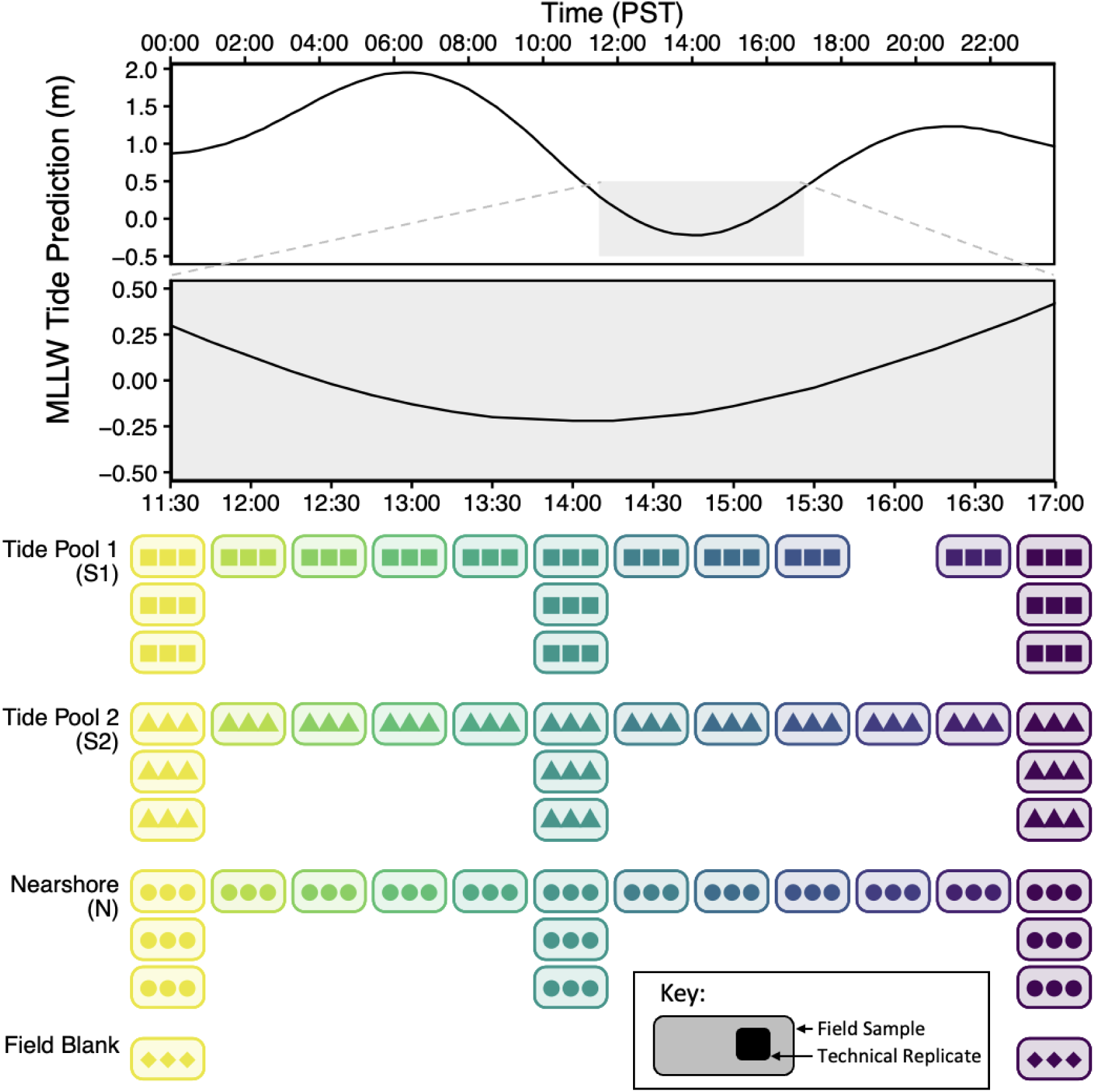
Graph of tide predictions during field sampling on 28 January 2022, with icons to indicate the sampling scheme.

At three time points (see Figure 3) at the beginning and end of the sampling period as well as at low tide (at 14:00 PST), we collected triplicate 1 L samples from each location as biological replicates. At the beginning and end of the sampling period, we also filtered 1 L MilliQ water via the procedure described above to serve as negative field controls. Additionally, using an Orion Model 1230 meter (Orion Research Inc., Beverly, MA, USA), we recorded temperature and salinity in each location directly after samples were collected.

Once finished filtering, Sterivex cartridges were dried by pushing air through them using a sterile 3 mL syringe, capped, placed in sterile Whirl-Pak bags (Whirl-Pak, Madison, WI, USA). Then, samples were stored in a cooler on ice until transported back to the laboratory at the end of the sampling period. Samples were transferred to a −20°C freezer for up to 18 days, at which time they were processed to extract nucleic acids from the captured materials.

### 2.4. DNA Extraction & Library Preparation

Within 18 days of collection, we extracted and purified DNA from the Sterivex cartridge using the DNeasy Blood and Tissue Kit (Qiagen, Germantown, MD, USA) and the modifications described in Spens et al. (2017). In short, we incubated the filter cartridge overnight with proteinase K and ATL. Then, we extracted the liquid from the cartridge with a syringe and mixed it with equal volumes of AL buffer and 0°C ethanol before proceeding with the manufacturer’s extraction protocol. One negative extraction control (DNA-grade water in place of a sample) was included in each of four batches of extractions. Nucleic-acids were stored at −20°C for up to 6 months.

We PCR amplified the extracted DNA in triplicate within 6 months of storage, using the mlCOIintF/ jgHCO2198 primer set targeting a 313 bp fragment of the mitochondrial COI region optimized by Leray et al. (2013) (Forward: GGWACWGGWTGAACWGTWTAYCCYCC; Reverse: TAIACYTCIGGRTGICCRAARAAYCA) with Nextera modifications. Following Curd et al. (2019), we used a 25 μl PCR reaction mixture consisting of 12.5 μl of Qiagen Multiplex Mix (Qiagen, Germantown, MD, USA), 2.5 μl each of the forward and reverse primers at 2 μM (Integrated DNA Technologies, Inc., Coralville, IA, USA), 6.5 μl of PCR-grade water, and 1 μl of undiluted DNA template. The PCR thermocycling touchdown profile began with an initial denaturation at 95° for 15 minutes to activate the DNA polymerase, followed by 13 cycles of denaturation (94° for 30 seconds), annealing (starting at 69.5° for 30 seconds, with the temperature decreased by 1.5° each cycle), and extension (72° for 1 minute). Then, an additional 35 cycles were run with the same denaturation and extension steps as above with an annealing temperature of 50°, followed by a final extension at 72° for 10 minutes. PCR reactions were prepared in a designated DNA-free hood until the template was added.

PCR amplification was conducted in two batches; in each batch, we included one no-template negative PCR control (DNA-grade water used as template). Additionally, we extracted and purified DNA from tissue from five organisms across a range of phyla we expected to amplify with the mlCOIintF/ jgHCO2198 primers, but not expected to be present at Pillar Point in particular (*Mytilus edulis, Mizuhopecten yessoensis, Xiphias gladius, Mercenaria mercenaria, Lutjanus campechanus*) using the standard tissue extraction protocol detailed in the DNeasy Blood and Tissue Kit (Qiagen, Germantown, MD, USA). These tissues were obtained from a local grocery store and it was assumed that they were labeled correctly, although previous work has indicated mislabeling in seafood stores can occur (e.g. Willette et al., 2017). Extracts from the 5 tissue samples were combined in equimolar amounts to form a mock community used as a positive PCR control in each batch. Triplicate PCR amplicons from both samples and controls were not subsequently pooled, but were carried through the remaining library preparation and sequencing steps as technical replicates. We electrophoresed and visualized a subset of PCR products on a 1.5% agarose gel stained with GelRed® (Biotium, Fremont, CA, USA) to ensure successful amplification and correct product sizes as well as lack of contamination.

Post-PCR library preparation and sequencing was conducted at the Georgia Genomics and Bioinformatics Core (GGBC, UG Athens, GA, RRID:SCR_010994). In short, provided PCR amplicons were cleaned using AMPure XP magnetic beads (Beckman Coulter, Indianapolis, IN, USA), barcoded with Nextera adapters (Illumina, San Diego, CA, USA) during a second PCR (3 min at 95 °C; 15 cycles of 30 sec at 95 °C, 30 sec at 67 °C, and 30 sec at 72 °C; and 4 min at 72 °C), cleaned again using AMPureXP magnetic beads, and pooled in equimolar ratio. The resulting library was sequenced on a MiSeq PE 2×250bp (500 cycles) using Reagent Kit V2 with 25% PhiX spike-in (Illumina, San Diego, CA, USA). Given our technical replication of samples and controls, our final library included 6 negative field controls (1 at beginning and end of field sampling, amplified in triplicate), 12 negative extraction controls (1 in each of 4 extraction sets, amplified in triplicate), 2 positive PCR controls (1 in each of 2 amplification batches), 2 no-template negative PCR controls (1 in each of 2 amplification batches), and 159 samples (53 field samples, amplified in triplicate).

### 2.5. Bioinformatics

We processed sequencing data using the *Anacapa Toolkit*, which contains two core modules: one for quality control and ASV parsing, and one for classifying taxonomy (Curd et al., 2019). Briefly, we ran the first module using default parameters, which uses *cutadapt* (version 1.16) (Martin, 2011) for adapter and primer trimming, *FastX-Toolkit* (version: 0.0.13) (Gordon & Hannon, 2010) for quality trimming, and *dada2* (version 1.6) (Callahan et al., 2016) for assigning ASVs. For the second module, we utilized the MIDORI2 reference database, a quality controlled and updated database built from GenBank release 253 (20 December 2022) that has been technically validated (Leray et al., 2022). Following Gold et al. (2022), we adjusted the identity and query coverage to 95% (default: 80%) to account for the relative incompleteness of the broad COI reference database compared to more taxonomically-specific databases (Curd et al., 2019). The second module relies on *Bowtie 2* (version 2.3.5) (Langmead & Salzberg, 2012) and a modified instance of *BLCA* (Gao et al., 2017) as dependencies. Following Gold et al. (2021) we only kept taxonomic assignments that had a bootstrap confidence cutoff score of 60 or higher in BLCA, to avoid spurious assignments from the incomplete reference database. We modified the Anacapa Container (Ogden, 2018), a Singularity container with all the needed dependencies for executing the Anacapa Toolkit, to enable the pipeline to be run in a high-performance computing environment requiring two-step authentication; the updated container, scripts, and reference database with the required *Bowtie 2* index library needed to reproduce our bioinformatics process are archived on Zenodo (Shea & Boehm, 2023a).

Raw ASVs, taxonomy assignments, and sample information were converted into interchangeable *ampvis2* (version 2.8.1) (Andersen et al., 2018) and *phyloseq* (version 1.44.0) (McMurdie & Holmes, 2013) objects in R (version 4.3.1) to facilitate decontamination and further analyses. Singletons were removed using *ampvis2*, and samples were further decontaminated using *phyloseq* by removing all ASVs that appeared in any negative field control, negative extraction control, or no-template negative PCR control, a choice made due to the very low number of overlapping ASVs between samples and negative controls (Table S1). Samples were rarified to the minimum number of reads of any sample using *ampvis2*. None of these decontamination steps changed the interpretation of subsequent analyses, as verified by replicating all analyses with datasets representing all 8 combinations of the presence/absence of the three decontamination and processing steps.

To ensure the accuracy of taxonomic assignments, we analyzed occurrence data from the Global Biodiversity Information Facility to investigate whether identified species had occurrence records in the California Current System and known ranges that encompassed Pillar Point, using the *spocc* (version 1.2.2) (S. Chamberlain, 2021) package. A phylogenetic tree based on taxonomic assignments was created using the *taxize* (version 0.9.100) (S. A. Chamberlain & Szöcs, 2013) and *ggtreeExtra* (version 1.10.0) (Xu et al., 2021) packages.

### 2.6. Data Analysis

To investigate whether eDNA signals could be distinguished by location, we first analyzed individual-level differences between locations; that is, whether ASVs or individual taxa (agglomerated species-level taxonomic assignments) were unique to, or associated with, particular locations. We calculated and visualized unique ASVs and taxa using the *eulerr* (version 7.0.0) (Larsson, 2022) package. However, with metabarcoding data in particular, taxa or ASVs that are unique to a given location are not necessarily ecologically meaningful; they could include rare taxa present elsewhere but not amplified and exclude taxa that are well correlated with particular locations but sometimes detected at others. Thus, we also analyzed ASVs and taxa using an indicator species framework (Dufrêne & Legendre, 1997). In this framework, the null hypothesis is that the frequency of taxon or ASV presence in samples from a particular location is not higher than the frequency of that taxon or ASV presence in samples from other locations. For each location, we identified all statistically-significant indicator taxa and ASVs with indicator value indices above 0.7—that is, indicators that are well-associated with a site group even if they are detected in samples from other sites—based on presence-absence data per sample using the *indicspecies* package (version 1.7.14) (Cáceres & Legendre, 2009).

We followed our individual-level analyses with approaches to test whether community composition varied by location. We calculated a Jaccard dissimilarity matrix across all samples using *vegan* (version 2.6.4) (Oksanen et al., 2022). Then, we tested for differences in community composition among locations using a permutational multivariate analysis of variance (PERMANOVA) with the model eDNA Presence ∼ Location + Time + Biological Replicates using the *adonis* function in *vegan*. We confirmed that locational differences found via PERMANOVA were not a result of differences in dispersions by testing for homogeneity of dispersions using the *betadisper* function in *vegan*. We also visualized Jaccard dissimilarity using non-metric multidimensional scaling (NMDS) using the *metaMDS* function in *vegan*. We coupled these analyses with a partitioning among medoids algorithm (Kaufman & Rousseeuw, 1990) implemented with the *pam* function in the *cluster* package (version 2.1.4) (Maechler et al., 2022). Rather than assuming location clusters a priori, we validated the optimal number of clusters for a given dataset by finding the number of clusters (k) that maximized the average silhouette width, a measure of how well-structured the clusters are. Finally, to better understand how the difference between locations varied over the time sampled, for each time point, we calculated the Jaccard dissimilarity between each unique combination of replicates within each site, and then the pairwise Jaccard dissimilarity between each unique combination of samples across the three pairs of sites: S1-N, S2-N, and S1-S2.

To further investigate whether differences in eDNA detections corresponded with underlying ecological gradients, we compared the unique and indicator taxa identified at each site to their ecological zonation in a highly regarded field guide to the Pacific intertidal, *Between Pacific Tides* (Ricketts et al., 1985). To account for potential variations in taxonomic names between *Between Pacific Tides* and the MIDORI2 reference database, we used the World Register of Marine Species (WoRMS) to identify all synonymized names for each unique and indicator taxon identified, and we searched *Between Pacific Tides* for all synonyms. We compared the proportion of high and middle intertidal species identified across each location using a chi-squared test with p-values computed via Monte Carlo simulation.

## 3. Results

### 3.1. Sequencing Results & Taxonomic Diversity

Using the Anacapa Toolkit, we identified 71,947 ASVs from 9,291,419 reads across 159 samples and 22 controls (positive PCR controls, negative field controls, negative extraction controls, and no-template negative PCR controls). All expected taxa were present in each positive PCR control, and no ASVs present in the positive PCR controls occurred in any other samples, providing no evidence of index hopping. Before application of any decontamination steps, 25 ASVs were shared between the samples and negative field, extraction, and PCR controls.

Focused on just the 159 samples, we removed singletons and all 25 ASVs that overlapped with negative controls (Table S1), and we rarefied samples to the minimum number of reads of any sample, leaving 27,038 ASVs and 4,047,981 reads. 89.1% of ASVs were taxonomically unassigned, and the remainder included representatives from 28 phyla, 59 classes, 132 orders, 234 families, 325 genera, and 415 species (Figure 4). A breakdown of the relative proportions of reads and ASVs assigned to each phylum can be found in Table 1. We confirmed that 291 of 415 identified species (70.1%) have occurrence records in the California Current System (CCS) and known ranges that encompass Pillar Point (additional details in SI 1 and Table S2).

**Figure 4:**
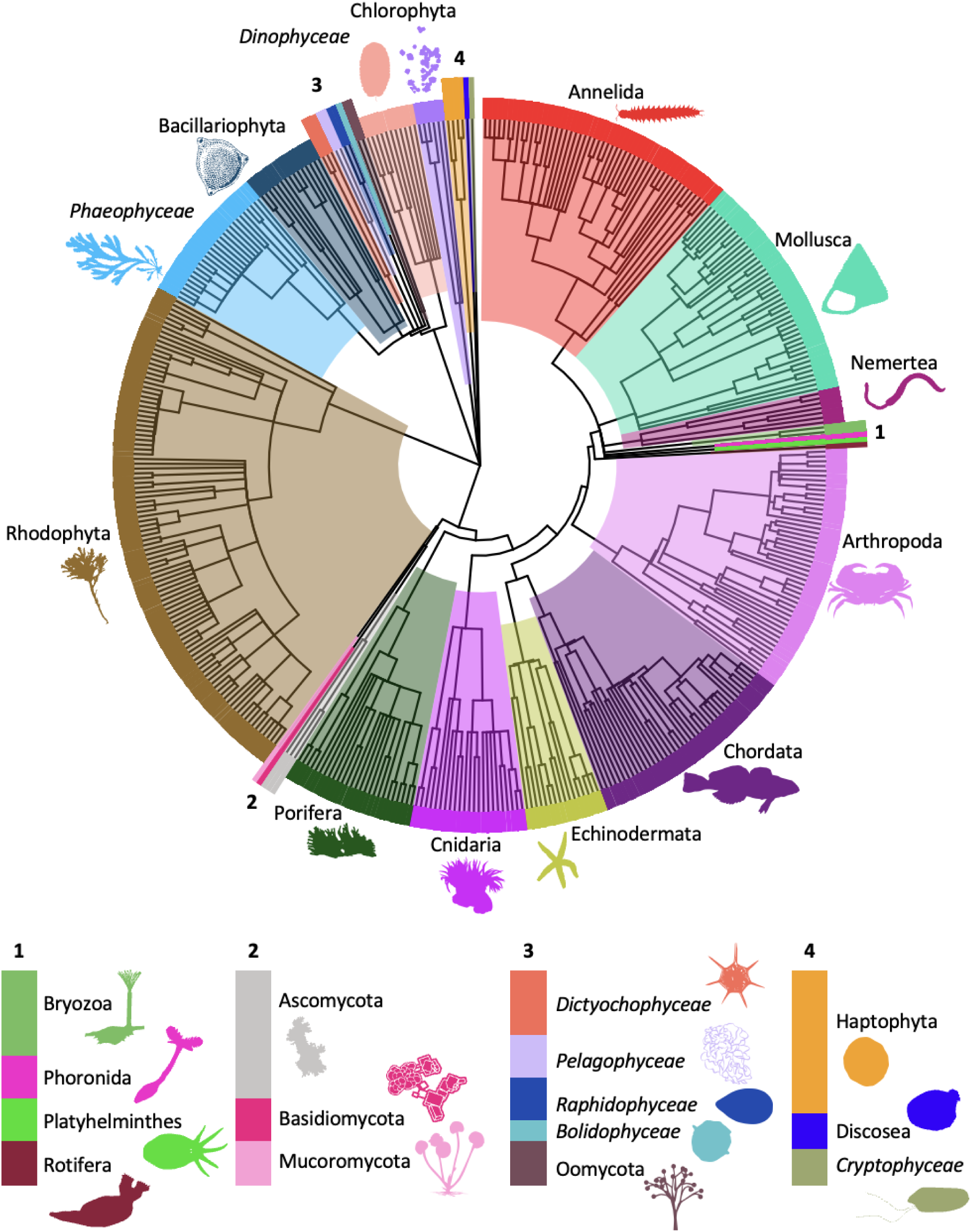
Phylogenetic tree colored by the phyla identified across all eDNA samples. Italicized taxonomic names are those with no phylum-level assignment, so class-level assignments are used to differentiate. Image Credit: PhyloPic.

**Table 1.** Chart of the total number, and relative proportion, of reads, ASVs, and species-level taxonomic assignments for each phylum. Italicized taxonomic names are those with no phylum-level assignment, so class-level assignments are used to differentiate.

|  | Reads |  | ASVs |  | Species-Level Assignments |  |
| --- | --- | --- | --- | --- | --- | --- |
| Phyla | Number | Percentage of Total | Number | Percentage of Total | Number | Percentage of Total |
| Rhodophyta | 532203 | 36.8% | 615 | 20.8% | 97 | 23.4% |
| Chlorophyta | 276017 | 19.1% | 189 | 6.4% | 6 | 1.5% |
| *Class: Dinophyceae* | 169579 | 11.7% | 120 | 4.1% | 8 | 1.9% |
| Arthropoda | 136328 | 9.4% | 423 | 14.3% | 48 | 11.6% |
| Chordata | 81310 | 5.6% | 173 | 5.9% | 39 | 9.4% |
| Annelida | 74621 | 5.2% | 477 | 16.2% | 50 | 12.1% |
| Cnidaria | 62818 | 4.3% | 153 | 5.2% | 22 | 5.3% |
| *Class: Phaeophyceae* | 42314 | 2.9% | 144 | 4.9% | 24 | 5.8% |
| Mollusca | 28102 | 1.9% | 267 | 9.1% | 43 | 10.4% |
| Echinodermata | 13406 | 0.9% | 90 | 3.1% | 14 | 3.4% |
| Porifera | 12900 | 0.9% | 114 | 3.9% | 20 | 4.8% |
| *Class: Dictyochophyceae* | 4030 | 0.3% | 10 | 0.3% | 3 | 0.7% |
| Haptophyta | 4019 | 0.3% | 27 | 0.9% | 2 | 0.5% |
| *Class: Pelagophyceae* | 3475 | 0.2% | 20 | 0.7% | 2 | 0.5% |
| Nemertea | 3095 | 0.2% | 23 | 0.8% | 7 | 1.7% |
| Bacillariophyta | 2101 | 0.1% | 51 | 1.7% | 15 | 3.6% |
| *Class: Bolidophyceae* | 453 | 0.03% | 3 | 0.1% | 1 | 0.2% |
| Phoronida | 110 | 0.008% | 12 | 0.4% | 1 | 0.2% |
| Bryozoa | 83 | 0.006% | 12 | 0.4% | 2 | 0.5% |
| Oomycota | 55 | 0.004% | 8 | 0.3% | 1 | 0.2% |
| *Class: Raphidophyceae* | 28 | 0.002% | 4 | 0.1% | 2 | 0.5% |
| Basidiomycota | 10 | 0.0007% | 2 | 0.07% | 1 | 0.2% |
| Platyhelminthes | 10 | 0.0007% | 4 | 0.1% | 1 | 0.2% |
| Ascomycota | 7 | 0.0005% | 4 | 0.1% | 3 | 0.7% |
| Discosea | 2 | 0.0001% | 1 | 0.03% | 0 | 0.00% |
| Mucoromycota | 2 | 0.0001% | 2 | 0.07% | 1 | 0.2% |
| *Class: Cryptophyceae* | 2 | 0.0001% | 1 | 0.03% | 1 | 0.2% |
| Rotifera | 2 | 0.0001% | 1 | 0.03% | 1 | 0.2% |
| **Total:** | **1447082** |  | **2950** |  | **415** |  |

### 3.2. Individual-Level Differences Between Locations

Across both taxa (n=415) and ASVs (n=27,038), all three locations had unique elements, suggesting that there was not complete mixing of eDNA across micro-habitats. 33.3% of taxa and 79.1% of ASVs were unique to one of the three locations (Figure 5). At both the taxa and ASV level, S1 had more unique elements than S2 and nearshore, although this effect was more pronounced across ASVs. ASVs and taxa shared across pairs of two locations were the least common elements.

**Figure 5:**
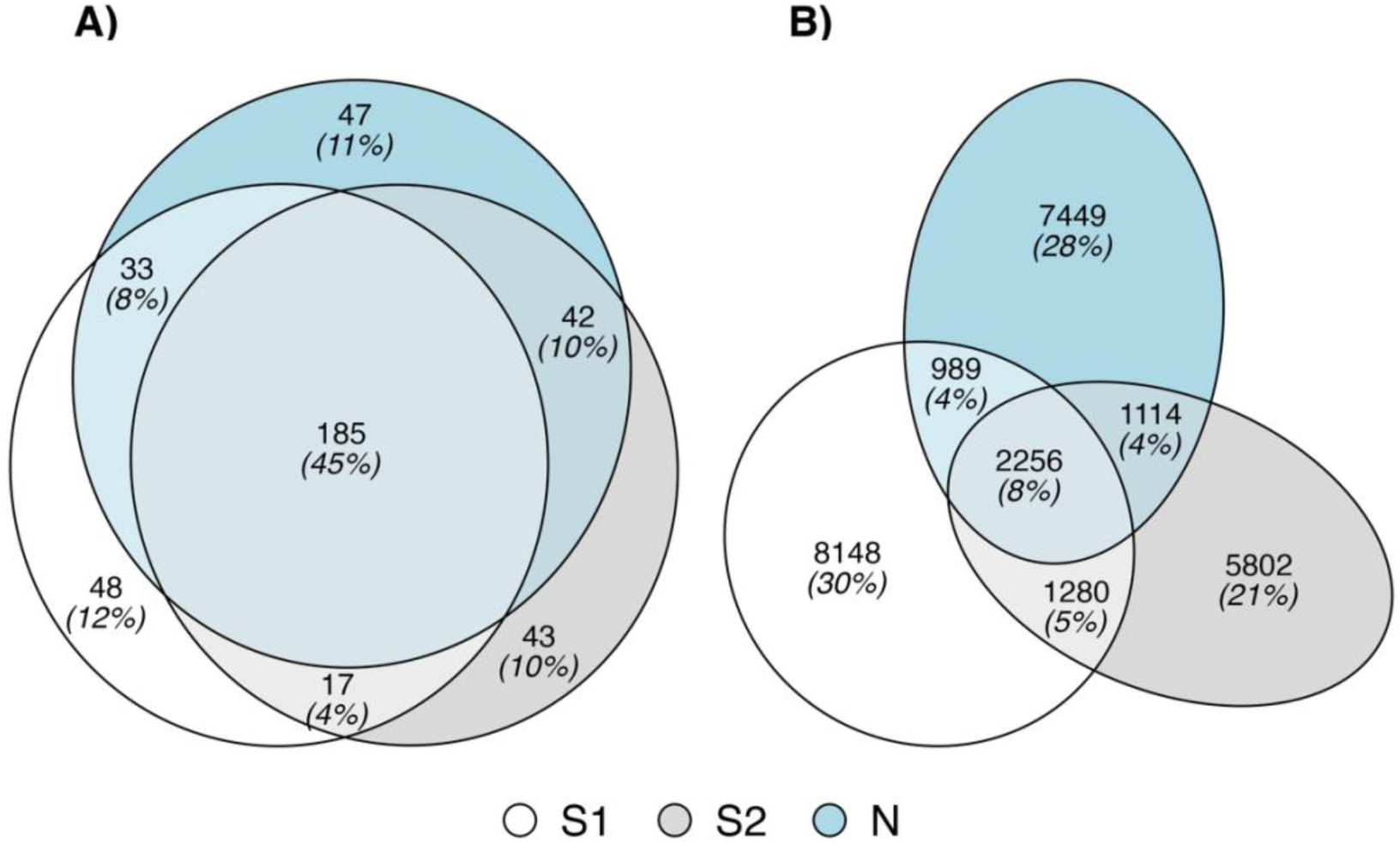
Area-proportional Euler diagrams showing the overlap in taxa (A) and ASVs (B) between locations S1, S2, and N.

Using an indicator species framework produced similar trends to the analysis of unique taxa and ASVs; all three locations had indicator taxa and ASVs, and S1 had more indicator elements than other locations. By taxa, we identified 11 indicator taxa: 7 for S1, 3 for S2, and 1 for nearshore (Figure 6). By ASVs, we found more indicators overall. We identified 99 indicator ASVs: 72 for S1, 10 for S2, and 17 for nearshore (Figure S1). Most (78.8%) indicator ASVs had no taxonomic assignment, but the ones that did included several species beyond the original indicator taxa, at S1 (*Bossiella frondescens, Bossiella plumosa, Corallina confusa, Leathesia difformis, Myrionema balticum, Scorpaenichthys marmoratus*), S2 (*Bossiella frondescens, Smithora naiadum*), and nearshore (*Halichondria panicea, Mazzaella splendens, Smithora naiadum, Strongylocentrotus purpuratus*).

**Figure 6.**
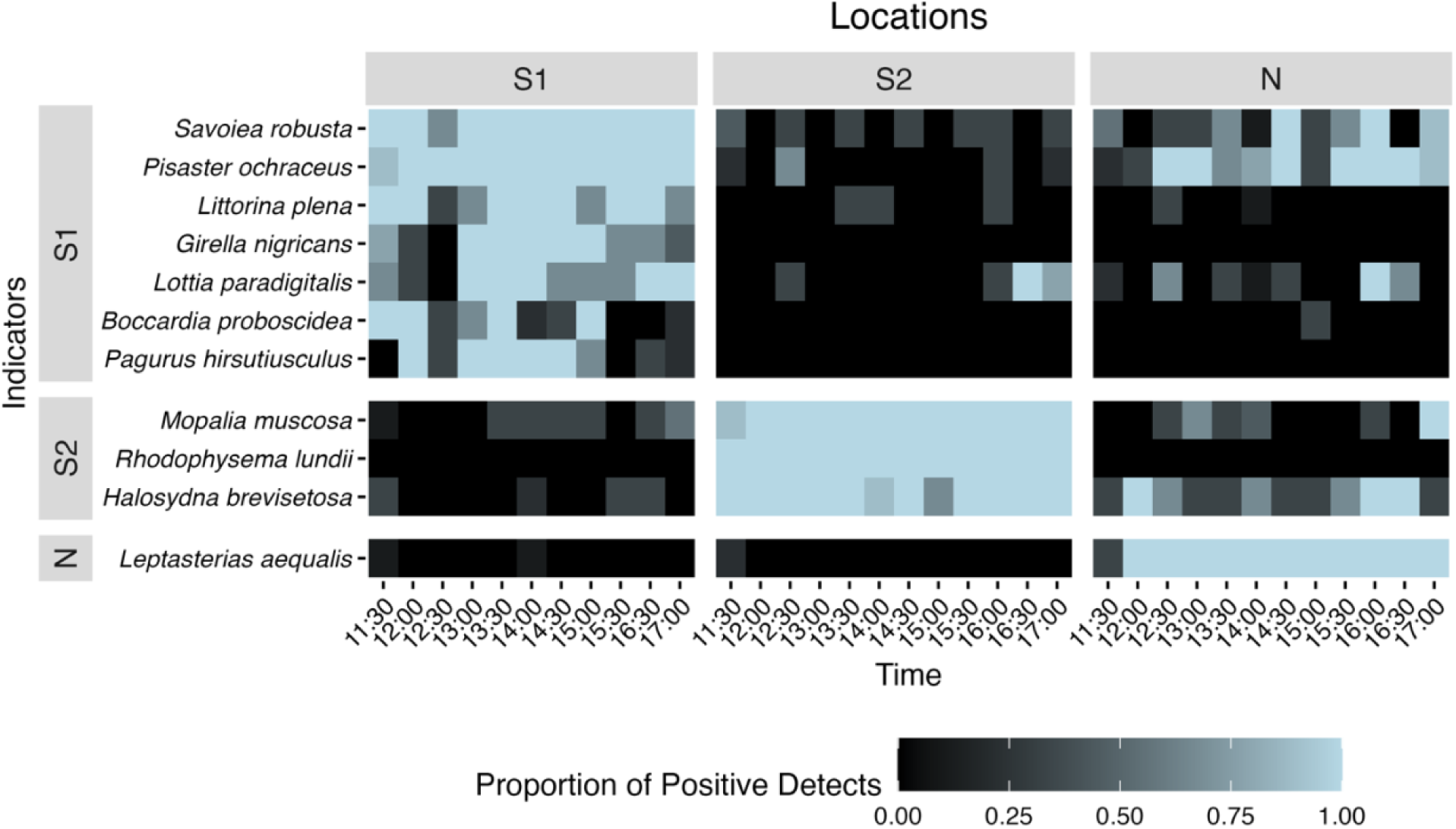
Heat map of identified indicator taxa for all three locations over time, scaled by the number of replicates that detected that species.

### 3.3. Community-Level Differences between Locations

Community composition also differed across locations, as supported by multiple analyses. Variation in community composition was significantly explained by location, even when accounting for variation due to time of sampling and variation between replicates (ASVs: PERMANOVA p < 0.001, betadisper p > 0.05; Taxa: PERMANOVA p < 0.001, betadisper p > 0.05). As visualized using NMDS ordination, S1 and S2 samples collected around low tide were most dissimilar from the offshore samples (Figures 7A, 7C). These patterns persisted in the optimal clusters identified using the partitioning among medoids (PAM) algorithm, without assuming a priori that location drives the clusters (Figure 7B, 7D). When analyzed by ASV, we identified three optimal clusters via PAM, primarily composed of: nearshore samples with additional samples from S1 and S2 at early time points (Figure 7B, Cluster 3), the remaining S1 samples (Figure 7B, Cluster 1), and the remaining S2 samples (Figure 7B, Cluster 2). When analyzing by taxa, we additionally differentiated between samples from the three locations collected at early time points via PAM (Figure 7D). Thus, multiple analyses with different initial assumptions all demonstrated differentiable eDNA signals across micro-habitats.

**Figure 7:**
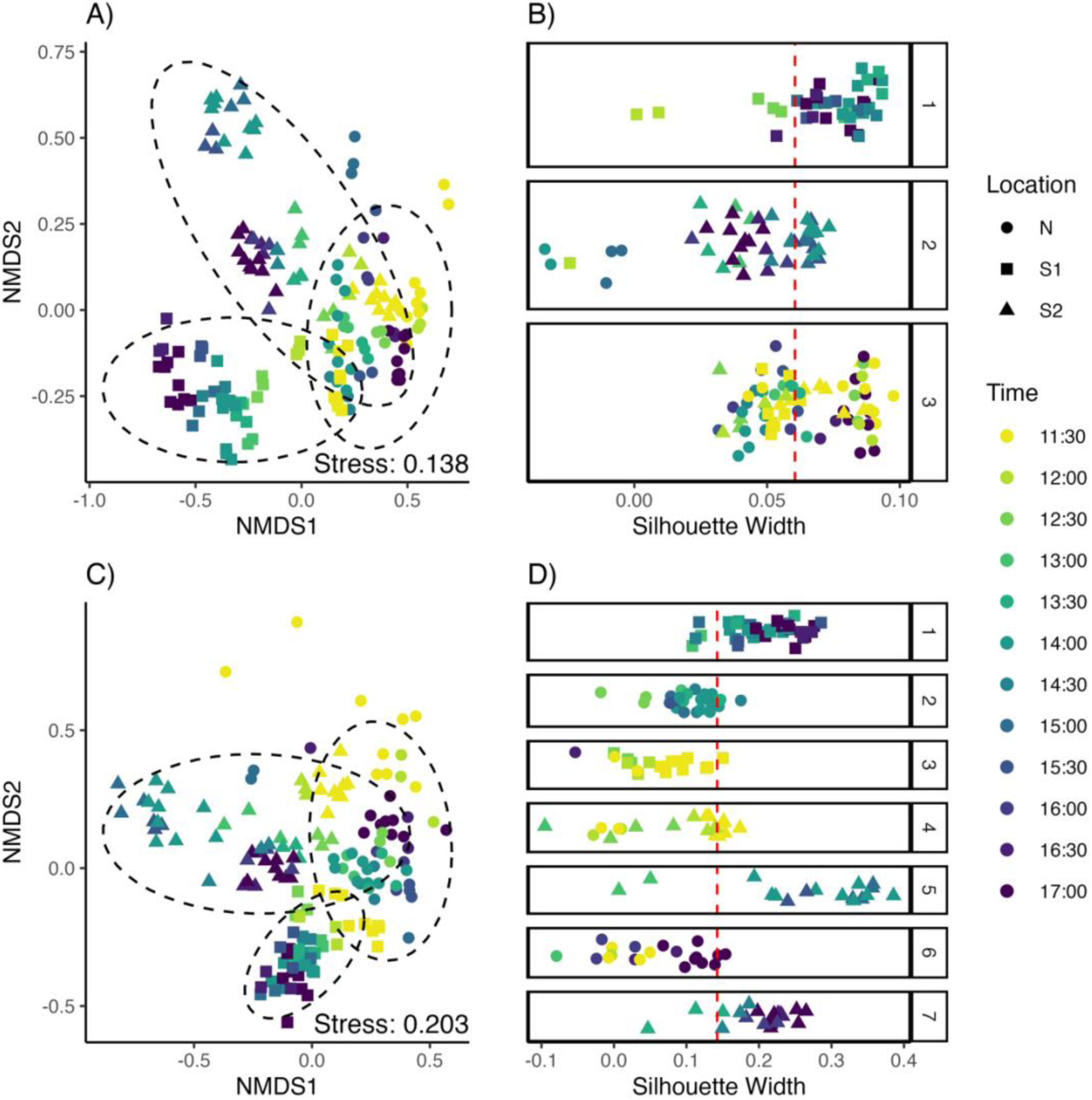
Ordination analysis (NMDS using Jaccard Distance; A, C) and cluster analysis (partitioning among medoids with number of clusters validated to maximize average silhouette width; B,D). Panels A & B are analyzed by ASV, and panels C & D are analyzed by taxa. 95% confidence ellipses for the centroids of each location based on a multivariate t-distribution are shown in the NMDS ordinations with black dashed lines, and average silhouette width is shown in cluster diagrams with a red dashed line. Points with larger silhouette widths represent elements more similar to their assigned cluster than other clusters.

The extent of the Jaccard dissimilarity between samples from different locations varied over time. As shown in Figure 8, except at the first time point sampled, the dissimilarity between samples across sites was always greater than the mean dissimilarity between samples within the same site, and increasingly so across the period sampled. At two time points—15:30 and 16:00—after low tide but before water was moving between all locations again, all dissimilarity values across sites were greater than the maximum dissimilarity recorded between two samples from the same site.

**Figure 8:**
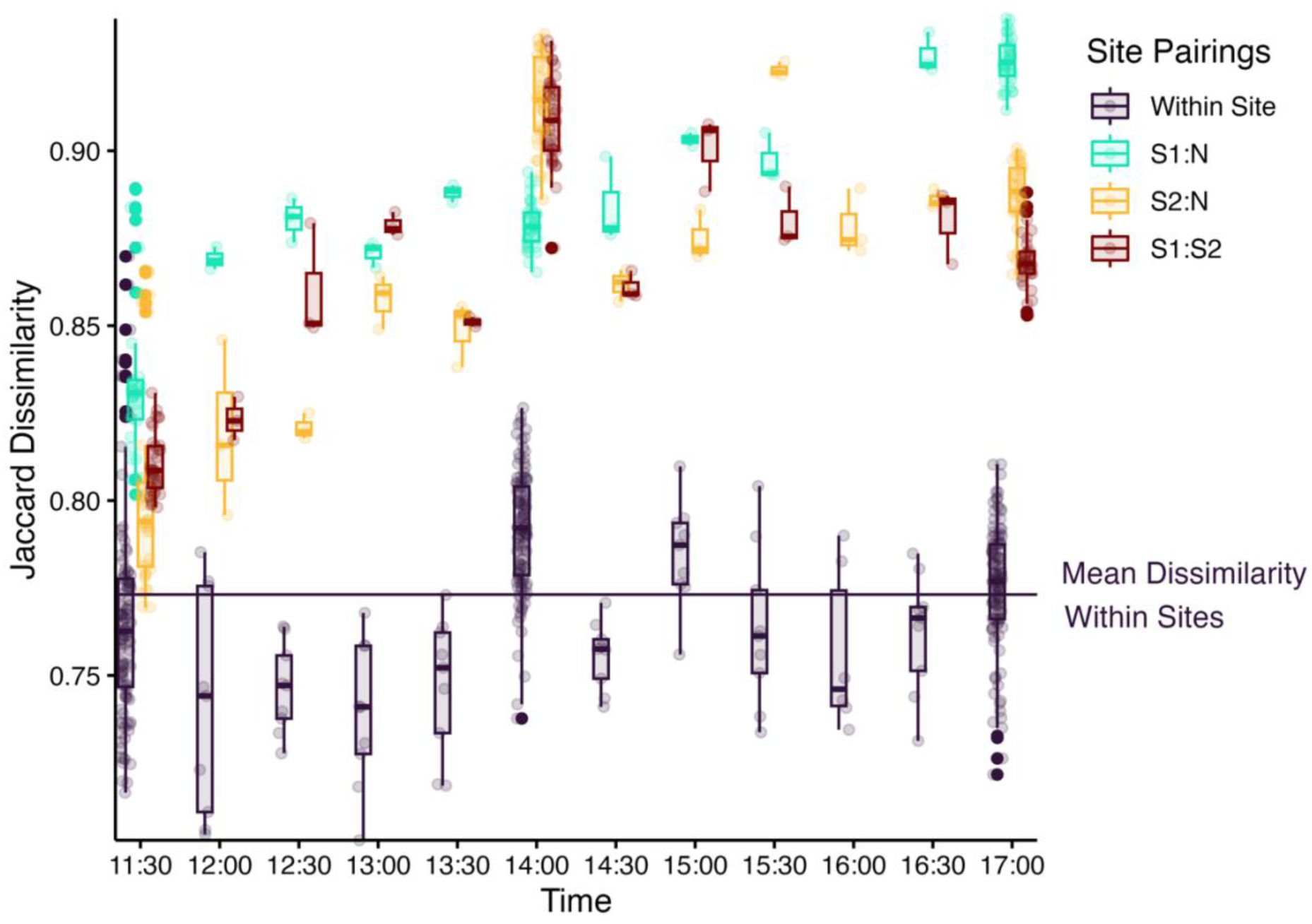
Box plots over time showing the pairwise Jaccard dissimilarity values between all unique pairs of replicates from the same sites (Within Site) and all unique pairs of samples across two sites (S1:O, S:O, S1:S2). The solid line depicts the mean Jaccard dissimilarity value from within site pairings.

### 3.4. Ecological Significance

Identified unique and indicator taxa reflected known ecological differences between locations. Only a subset of unique and indicator taxa were described in *Between Pacific Tides*: 18.8% (9/48) of nearshore taxa, 32.1% (17/53) of S1 taxa, and 31.1% (14/45) of S2. However, across the subset present in *Between Pacific Tides*, more taxa from S1 were categorized to high and middle intertidal zones than S2 and N (Figure 9). The proportion of taxa from the high and middle intertidal varied significantly across locations (*χ*2 = 11.67, *p* < 0.05), matching the environmental characteristics of the sites.

**Figure 9.**
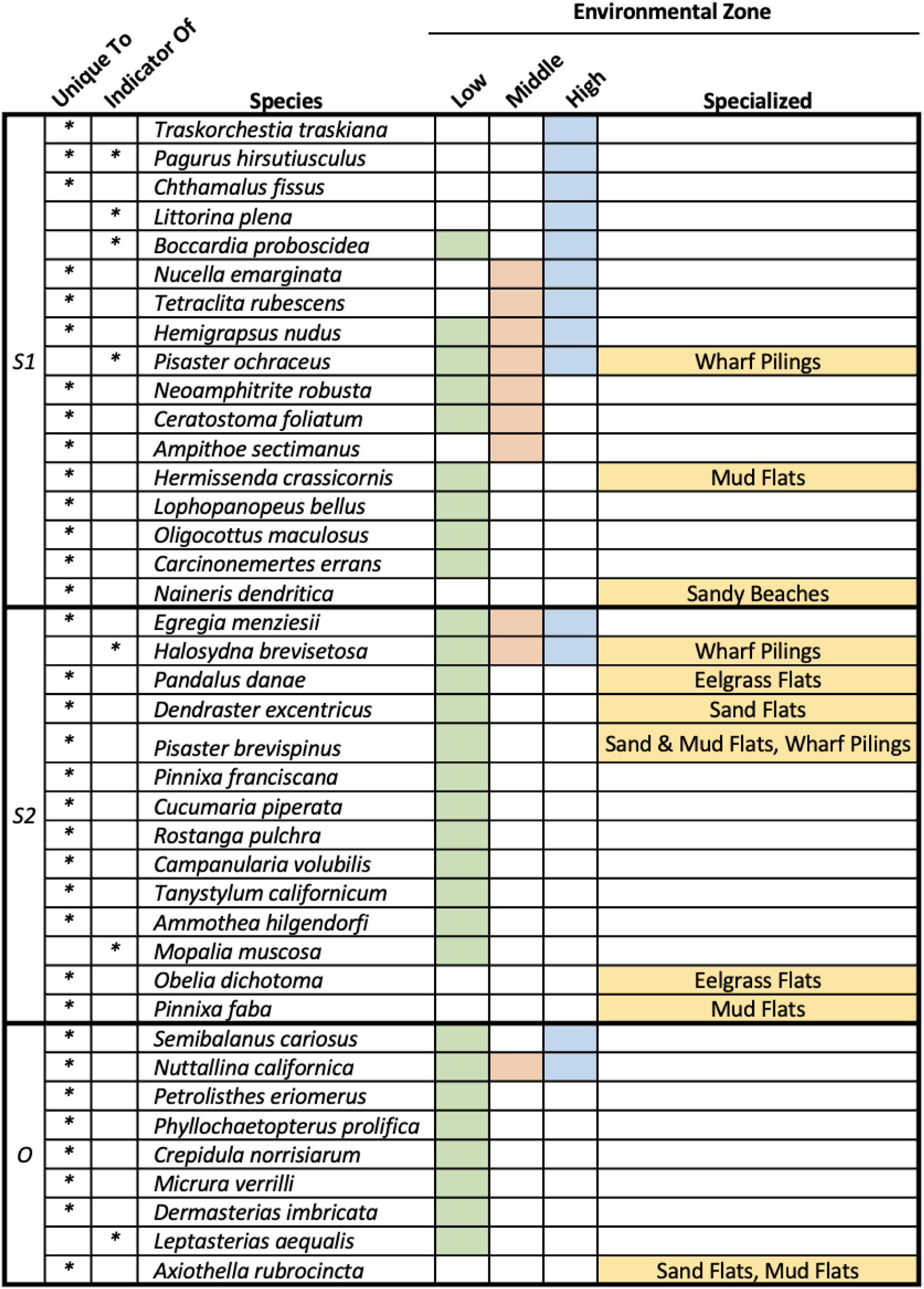
Chart habitat information about unique and indicator species present in Between Pacific Tides.

## 4. Discussion

### Differentiation between rocky intertidal micro-habitats via eDNA metabarcoding survey

We detected different eDNA signals across rocky intertidal micro-habitats separated by approximately 40 meters, both when analyzing individual differences in ASVs and taxa and when analyzing multivariate community composition. The eDNA composition of samples from S1 and S2 were both generally more different from nearshore samples after low tide than before low tide, supporting the notion that the degree of physical connectivity influences the detectability of a unique eDNA signal from micro-habitats. However, we did not observe a direct correspondence between the physical connectivity of our locations and the strength of their eDNA signal. At both the start and the end of the sampling period, water was actively flowing between all three locations. However, the three locations remained more differentiable from one another at the end of the sampling period than the beginning, despite being similarly physically interconnected. Consistent with prior work showing no significant tidally-driven flux of offshore eDNA (Kelly et al., 2018), this may indicate that when sites are fully isolated for a period, allowing localized shedding to drive the eDNA signal, the strength of that signal is resilient to moderate reintroduction of transport.

These findings point to the need for even finer-grained analyses of spatiotemporal variability of eDNA in complex coastal environments. We selected our two tide pool locations because they were strictly isolated for some portion of their exposure period and represented quite different ecological zones in the intertidal despite their proximity. But the strength of their differentiability suggests that even finer-scale differences might be possible to detect, and that full physical isolation may not be necessary to enable those detections. Continued work at smaller spatial scales, and at additional locations, will help bolster understandings of the limit of differentiability between sites using eDNA metabarcoding in complex coastal environments. Furthermore, monitoring beyond the points in the tidal cycle where locations are physically accessible by foot—the core factor limiting our sampling period—will help resolve the extent to which unique eDNA signals can persist even with mixing.

### Concordance with ecological expectations

Beyond just detecting differences between micro-habitats, we found that those differences matched ecological expectations. Site S1 was located closer to shore and interior to a channel that divides the Pillar Point intertidal, characteristic of the high to middle intertidal. Site S2 was located across the channel and further from shore, characteristic of the low intertidal. Given this, and despite their equidistance from the nearshore sampling location, we expected that the higher-zoned S1 would be more different from the nearshore samples than the low-zoned S2. Additionally, we expected that S1 would contain more unique and indicator species known to be characteristic of the high and middle intertidal.

Our analyses confirmed both expectations. Across most time points, samples from S1 were more different overall from nearshore samples than were samples from S2. This trend was paralleled in individual-level analyses as well; S1 had more unique species and taxa than S2, as well as more indicator species and taxa. Furthermore, across unique and indicator species identified, more of the species from S1 were representative of the high and middle intertidal than those from S2.

These ecological comparisons are necessarily limited. Significant prior research has compared eDNA metabarcoding surveys to visual surveys in marine environments (e.g. Gold, Sprague, et al., 2021; Kelly et al., 2017; Port et al., 2016), including early work in the intertidal in particular (R. S. Meyer et al., 2019). Meta-analyses have shown that generally, eDNA metabarcoding surveys are comparable to visual surveys (Fediajevaite et al., 2021; Keck et al., 2022; McElroy et al., 2020), although the particular complementarity between methods varies by context and research approach. Thus, our aim in this study was not to directly compare eDNA metabarcoding with conventional intertidal surveys, so our ecological inferences were based on general natural history observations rather than a direct comparison with visual observations. Future work directly comparing intertidal surveys with eDNA surveys could enhance ecological interpretation of eDNA results from rocky intertidal ecosystems. But even absent additional field sampling, exploring networks of associations between identified taxa and ASVs (e.g. Djurhuus et al., 2020; R. S. Meyer et al., 2019) could enable further ecological analyses. As a first step, we have motivated the utility of this future work by demonstrating that eDNA surveys can indeed resolve small spatiotemporal differences in rocky intertidal ecosystems, and that those differences are ecologically meaningful.

### Limitations and variability in eDNA metabarcoding surveys

While our results highlight the promise of eDNA metabarcoding for detecting small-scale ecological differences, they also demonstrate some of the ongoing challenges in adopting eDNA analyses for routine biomonitoring. For one, we found that even biological and technical replicates were highly dissimilar, consistent with other eDNA metabarcoding studies (e.g. Beentjes et al., 2019; Shirazi et al., 2021; Stauffer et al., 2021). That micro-habitats were still differentiable despite variability across replicates is promising, and suggests that observed differences between micro-habitats were driven by differences in core ecological communities rather than rare taxa. At the same time, it demonstrates that additional replication would be needed to more exhaustively survey these micro-habitats (Stauffer et al., 2021).

Additionally, like many eDNA metabarcoding studies utilizing “universal” COI assays (e.g. Jeunen et al., 2019), a high proportion of ASVs could not be taxonomically assigned; 88% of ASVs from this study could not be assigned to species. Many factors contribute to the limitations of COI assays, from incomplete reference databases (Porter & Hajibabaei, 2018; Weigand et al., 2019) to a lack of conserved primer binding sites within the COI protein-coding gene that can result in unreliable amplification across broad taxonomic ranges (Deagle et al., 2014). Additionally, conservative choices made in our bioinformatics processing (e.g. our high threshold for identity and query coverage) prioritized accuracy in taxonomic assignments, but at the cost of a higher proportion of unassigned ASVs. At a high level, these trade-offs and shortcomings did not impact the interpretation of our results; both our ASV and taxa-level analyses showed differences among micro-habitats. However, when trying to make ecological sense of those differences, a large portion of our dataset without taxonomic assignments was excluded. Well-established approaches exist for improving biodiversity coverage from eDNA metabarcoding surveys, including constructing local reference databases to ensure complete coverage of taxa of interest (Gold, Curd, et al., 2021) and using multiple metabarcoding assays in tangent (Kelly et al., 2017; Stat et al., 2017). Additionally, there is a growing interest in biological interpretation using taxonomy-free eDNA approaches (Apothéloz-Perret-Gentil et al., 2017). However, these approaches are not without shortcomings; the taxonomy-based fixes are resource intensive, and the taxonomy-free approach can only address some biomonitoring priorities.

Furthermore, while we saw similar high-level trends in our ASV and taxa level analyses, the degree to which these different analytical levels could resolve differences between micro-habitats varied across our analyses. We found a higher proportion of unique ASVs than unique taxa, and differences in Jaccard dissimilarity over time were greater when analyzing by ASV than analyzing by taxa. However, PAM clustering was better able to differentiate between micro-habitats using taxa than using ASVs. In our case, these variations in degree of differentiation did not influence our interpretation: across all ASV and taxa level analyses, using a variety of multivariate and univariate approaches, we found evidence that eDNA metabarcoding could differentiate between distinct locations. However, that one analytical level (ASV vs. taxa) was not consistently a stronger determinant than the other indicates that for other research questions and contexts, different analytical approaches could yield conflicting results. Given the variability and limitations of eDNA metabarcoding surveys, and especially the data loss when analyzing by taxonomy, our results emphasize the value of combining different analytical approaches to increase confidence in ecological findings.

### Pragmatic Implications for Rocky Intertidal Biomonitoring

Many researchers have highlighted the importance of pilot sampling to understand localized spatial and temporal eDNA variability, and thus to determine the best sampling plan for answering the research question(s) of interest (e.g. Bruce et al., 2021; Gold et al., 2022). Pragmatically, our findings support this need. Rocky intertidal ecosystems may represent particularly practical locations for establishing routine eDNA biomonitoring. They can be more easily accessible than other commonly sampled coastal locations (e.g. surf zones), potentially making routine sampling more feasible. Additionally, they have been important sentinel sites for studying the impact of global environment changes (e.g. Barry et al., 1995; Helmuth et al., 2006; Mieszkowska, 2021; Sagarin et al., 1999). Furthermore, intertidal sites are already common locations for community science monitoring (Thiel et al., 2014) and popular recreational destinations, potentially enabling increased eDNA sample collection by the public (e.g. R. Meyer et al., 2021). Yet, our findings demonstrate that different sampling schemes in the rocky intertidal could potentially inform very different biomonitoring questions. Sampling in one individual tide pool right when it is accessible by foot might help generate a broad biodiversity signal for the full ecosystem, but sampling in the same pool after it has been physically isolated might help uncover the specific characteristics of that micro-habitat. Future work in additional intertidal locations will be needed to understand if there are generalizable trends in rocky intertidal eDNA dynamics that could inform sampling best practices. Our findings simply showcase the wide range of research questions that may be possible to answer by collecting eDNA samples in the rocky intertidal, and the importance of considering physical connectivity when determining the best sampling plan to answer those questions.

## 5. Conclusion

Monitoring biodiversity changes, especially in marine environments, remains challenging, resource-intensive, and crucial for continuing efforts to understand and mitigate human-induced environmental impacts. While the utility of eDNA biomonitoring has been broadly demonstrated, important questions remain about its spatiotemporal variability, potentially complicating the introduction of eDNA biomonitoring to new ecosystems and research contexts. Our results demonstrate that eDNA biomonitoring can differentiate micro-habitats in the rocky intertidal only 40 m apart, that these differences are ecologically meaningful, and that physical connectivity informs the degree of differentiation possible. This small-scale differentiability equips eDNA biomonitoring to contribute to many types of research questions in rocky intertidal ecosystems: from tracking small scale changes in organism ranges to broad shifts in community composition.

Monitoring changes over small spatial scales in the intertidal has enabled some of the earliest characterization of climate impacts in marine ecosystems (e.g. Barry et al., 1995; Sagarin et al., 1999), and remains an important mechanism for studying global environment changes (e.g. Mieszkowska, 2021). These conventional intertidal surveys will likely never be fully replaceable by eDNA biomonitoring. Translating an eDNA signal to abundance data (Shelton et al., 2023), characterizing the full extent of eDNA variability (Mathieu et al., 2020), and managing eDNA data (Shea et al., 2023) all remain challenges, among many others. Yet, our findings suggest that eDNA biomonitoring could be an important complement to existing intertidal monitoring efforts. Consistent with other studies demonstrating the small-scale differentiability of eDNA signals in coastal ecosystems (e.g. Gold, Sprague, et al., 2021; Port et al., 2016), our data demonstrates the potential power of eDNA biomonitoring to increase the spatial and temporal resolution of marine biodiversity data, with implications for research, conservation, and management.

## 6. Acknowledgements

Thank you to Callie Chappell, Noah Gluschankoff, Heidi Hirsh, Jacob Kuppermann, Tyler Leeds, Ryan OConnor, and Ryan Searcy for assisting with field sampling; David Mucciorone for lending field equipment; Eily Andruszkiewicz Allan, Zachary Gold, and Tanner Waters for guidance on field and laboratory methods; Tadashi Fukami, Jesse Miller, and Robin Elahi for guidance on statistical analyses; and Kelly Dunn for graphic design work on Figure 1. Sequencing data were generated at the Georgia Genomics and Bioinformatics Core (GGBC, UG Athens, GA, RRID:SCR_010994); we are grateful as well for their troubleshooting and logistical support. Some of the computing for this project was performed on the Sherlock cluster. We would like to thank Stanford University and the Stanford Research Computing Center, especially Zhiyong Zhang, for providing computational resources and support that contributed to these research results.

Funding was obtained from the Emmett Interdisciplinary Program in Environment & Resources and the Stanford Doerr School of Sustainability McGee-Levorsen Grant Program; thanks to Ann Marie Pettigrew, Ai Le Tran, Gabriela Magana, Alyssa Ferree, and Carine Sauquet for administrative support.

## 8. Data Accessibility and Benefit-Sharing

### Data Accessibility Statement

We have published detailed, step-by-step protocols for many of the methods described, including specific materials used, photographs, and additional methodological notes. See Shea and Boehm for sample collection and filtering (2023b), for DNA extractions (2023c), for PCR amplification (2023d), and for shipping samples (2023e).

Our main hub for data availability is on Dryad (Private Reviewer Link: https://datadryad.org/stash/share/Kxkudmvtnl8nEBBi3NBo2VFxFeluDLh1WlhmgR66lR0), which contains FASTQ files and eDNA datasets (pre-processing & post-processing). Our Dryad repository also contains links to the workflows and code needed to generate and analyze these datasets: our modified Anacapa Container and scripts for bioinformatics (Shea & Boehm, 2023a), as well as our GitHub repository including an R Markdown file that reproduces all methods & results detailed here (Shea & Boehm, 2023f).

Our FASTQ files will also be uploaded to the NCBI SRA (accession number forthcoming), and our processed dataset will be uploaded to GBIF (accession number forthcoming).

### Benefit-Sharing Statement

Benefits from this research accrue from the sharing of our protocols, data, and results on public databases as described above.

