## Supplementary figures and images for "Environmental DNA (eDNA) metabarcoding differentiates between micro-habitats within the rocky intertidal"

### Supplemental Figure S1

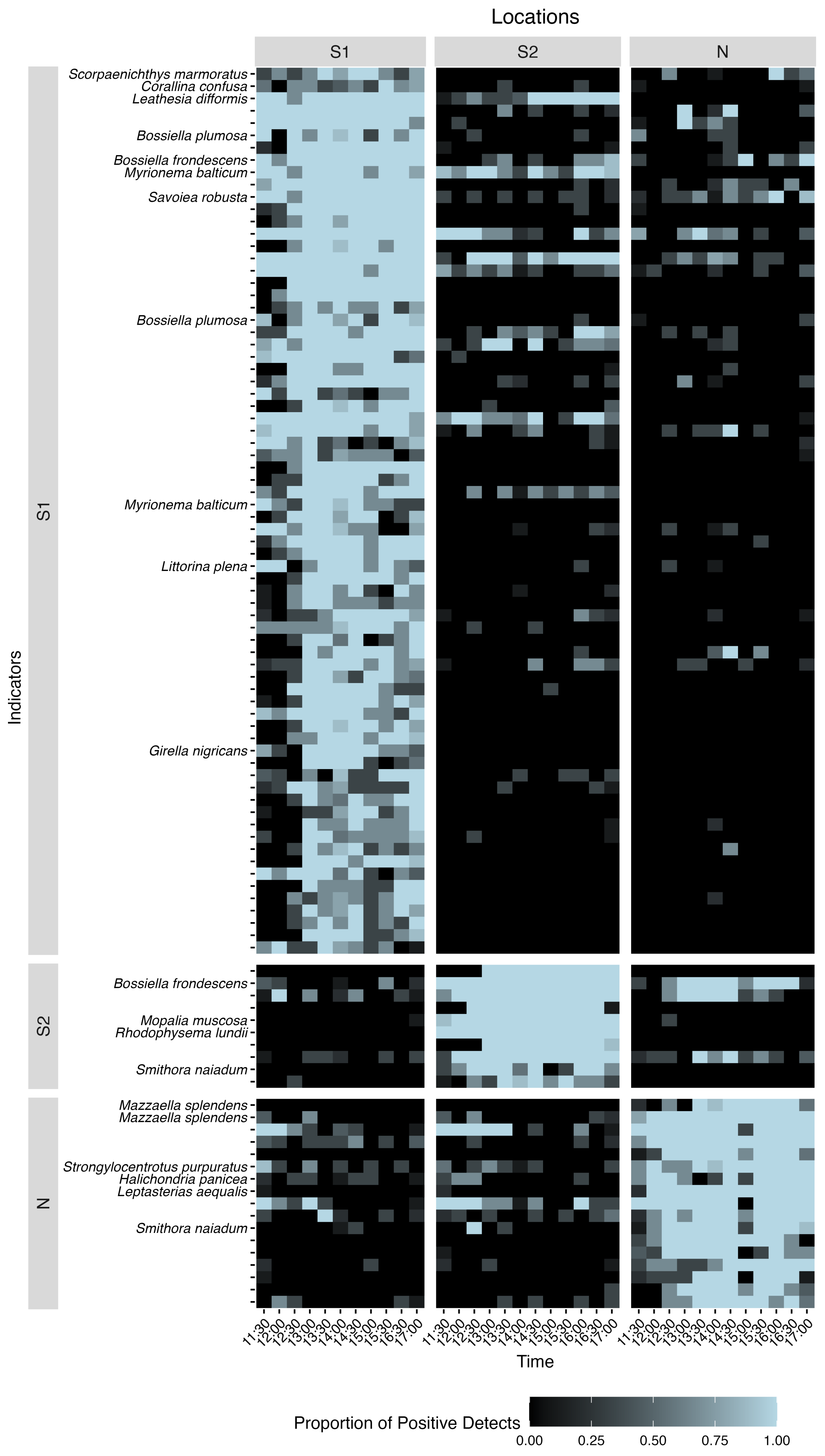
