## Supporting Information 1 for "Environmental DNA (eDNA) metabarcoding differentiates between micro-habitats within the rocky intertidal"

*SI 1: Full GBIF Analysis Details*

By analyzing occurrence data from the Global Biodiversity Information Facility, we confirmed that 100 of 415 identified species (24.1%) have occurrence records at Pillar Point, specifically. An additional 191 identified species (46.0%) have occurrence records in the California Current System (CCS) and known ranges that encompass Pillar Point. Of the remaining species, some only fulfilled one of two criteria; 12 identified species (2.9%) had occurrence records in the CCS but known ranges that did not encompass Pillar Point and 35 identified species (8.4%) had ranges that encompass Pillar Point but no occurrence records in the CCS. Additionally, 41 identified species (9.9%) had neither occurrence records in the CCS nor known ranges that encompassed Pillar Point. Finally, 16 identified species (3.9%) lacked sufficient occurrence records in GBIF to make any designation, and 20 identified species (4.8%) did not have species-level records in GBIF. Full details by species can be found in Table S2.
